# Small Heat Shock Proteins and Eicosanoid Pathways Modulate Caspase-1 Activity in the Fat Bodies of *Antheraea pernyi*

**DOI:** 10.1101/026039

**Authors:** Congfen Zhang, Lei Wang, Guoqing Wei, Cen Qian, Lishang Dai, Yu Sun, Baojian Zhu, Chaoliang Liu

## Abstract

After heat shock injury, a group of proteins that regulate protein-folding processes are synthesised to prevent damage. Caspase is an enzyme responsible for the execution ostress-induced apoptosis. Heat shock proteins (Hsp) are capable of modulating caspase activity. In addition to changes in protein synthesis, heat shock causes the release of arachidonic acid (AA) from plasma membranes and the subsequent synthesis of eicosanoids, i.e., activation of the AA pathways. The development of cytoprotective strategies might depend on whether caspase-1 activity is affected by heat shock preconditioning and the associated pharmacological modulations after heat shock injury. Therefore, we studied the effects of heat shock preconditioning and modulations of the eicosanoid pathways on *Ap-sHSP20.8* and on the final apoptotic effector enzyme caspase-1 to clarify whether these effects were modulated in the fat bodies of *Antheraea pernyi* injured by heat shock. We concluded that eicosanoid biosynthesis inhibitors might be utilised to simultaneously decrease Hsp 20.8 synthesis and to increase caspase-1 activity. Modifications of the eicosanoid pathways might also be used to mediate caspase-1 activity under hyperthermic conditions, suggesting a novel mechanism for regulation of caspase-1 in the fat bodies of *A. pernyi*

## 1. Introduction

Protein-damaging cellular stress, such as exposure of cells to elevated temperature, activates an adaptive response that increases the synthesis of a group of proteins that regulate protein-folding processes. An increase in cellular temperature causes denaturation of proteins and interrupts critical cellular processes, and thus, the result is apoptosis and cell death. Heat shock may induce apoptosis in many cell lines not only directly but also indirectly by activation of most apoptotic stimuli, including activation of acid sphingomyelinase(Haimovitz-Friedman et al, 1997) in preimplantation embryos and oocytes(Paula-Lopes and Hansen, 2002) and in other human and mouse cells (Ding, 2014; Marfe et al, 2009; Nagasawa et al, 2014; Shi et al, 2004). Apoptosis, or programmed cell death, is orchestrated by a family of proteases known as caspases that cleave their substrates after specific aspartic acid residues. At least 14 caspases are found in mammalian cells, which are synthesized as inactive precursor molecules (procaspases) and are converted by proteolytic cleavage to active heterodimers (Thornberry and Lazebnik, 1998). Lepidoptera possess at least 5 caspases(Courtiade, 2011; Huang, 2013). Heat shock proteins (HSPs) are well established as stress-induced molecular chaperones, which repair denatured proteins and confer protection against apoptotic stimulation (Dick et al, 2000). The overexpression of HSP 70 or HSP 27 makes cells resistant to death, and heat-induced apoptosis is blocked in Hsp70-expressing cells. This was associated with reduced cleavage of the common death substrate protein poly (ADP-ribose) polymerase (Dick et al, 1997). Additionally, the expression of some HSPs, such as the tissue-specific expression of Hsp70-2, prevented apoptosis in spermatocytes(Davidet al, 1996). Sorbitol exposure caused apoptosis in the K562, U937 and HeLa cell lines with a marked activation of caspase-9 and caspase-3. However, heat-shock pretreatment before sorbitol exposure induced the expression of Hsp70 and inhibited the sorbitol-mediated cytochrome c release and the subsequent activation of caspase-9 and caspase-3. Similarly, overexpression of HSP70 in three cell lines prevented caspase-9 cleavage and activation as well as cell death(Marfe et al, 2009). In the plantaris of adult rats, Hsp25 and pHsp25 were negatively correlated with caspase-3 activity, and Hsp25 was correlated with muscle mass(Huey et al, 2008). Additionally, yeast Hsp104 facilitated resolubilization of insoluble protein aggregates during stress (Parsell, 1994). By contrast, heat shock inhibition of active caspase-1 occurred independently of an inflammasome platform through a titratable factor present within intact, functioning heat-shocked cells. High temperatures significantly promoted the apoptosis process, which resulted in an increased mortality rate for *Plutella xylostella* (Zhuang, 2011). In Lepidoptera, apoptosis is essential in processes such as metamorphosis or defending against baculovirus infection. Caspase-1 is an enzyme responsible for the execution of stress-induced apoptosis. However, in the Chinese silk worm, *Antheraea pernyi*, it is presently unknown whether the HSP family shares similar mechanisms of cytoprotection.

In addition to changes in protein synthesis, heat shock induces the release of arachidonic acid (AA) from plasma membranes and the subsequent synthesis of eicosanoids, i.e., activation of the AA pathways. Eicosanoids are oxygenated metabolites of polyunsaturated fatty acids, particularly arachidonic acid. Eicosanoid chemical structures and biosynthetic pathways are detailed elsewhere(Stanley and Kim, 2011; Stanley, 2006). In insects, eicosanoids mediate cellular immunity to microbial and metazoan challenges(Stanley and Kim, 2014). Notably, some injured organisms secreted factors responsible for impairing host insect immune reactions by inhibiting the biosynthesis of eicosanoids (Buyukguzel et al, 2009). In mammalian cells, heat shock (42-45 °C) increased the rates of arachidonic acid release from human, rat, murine, and hamster cells(Calderwood et al,1989). Heat shock proteins (HSPs), including Hsp60, are capable of modulating caspase-3 activity. Heat shock preconditioning upregulates Hsp synthesis and inhibits restitution and cell proliferation via mechanisms related to de novo protein synthesis and eicosanoid pathways. These are both essential in the regulation of apoptosis and the gastric mucosal defence systems(Oksala et al, 2004).

The Chinese oak silk moth *Antheraea pernyi* (Lepidoptera: Saturniidae) is an economically valuable silk-producing insect that is commercially cultivated primarily in China, India, and Korea(Zhou and Han,2006). In this study, we studied the effect of heat shock preconditioning and modulations of the eicosanoid pathways on *Ap-sHSP20.8* and on the final apoptotic effector enzyme caspase-1 to clarify whether these effects were modulated in the fat bodies of *A. pernyi* injured by heat shock.

## 2. Materials and Methods

### 2.1. Experimental insects

The larvae of *Antheraea pernyi* were from the Sericultural Research Institute of Henan province, and the larvae were reared on oak leaves (Wei et al, 2006). Ten randomly selected fifth-instar larvae (the third day of the fifth-instar) were exposed to 43 °C for 0.5 h in a constant temperature incubator and then were kept at 25 °C for 1 h. After heat treatment, the fat bodies were immediately collected, frozen in liquid nitrogen and stored at -80 °C.

### 2.2. RNA extraction and cDNA synthesis

Total RNA was extracted from fat bodies with the EASYspin Total RNA Extraction Kit (Aidlab Biotech Co. Ltd., Beijing) according to the manufacturer’s instructions. 800 ng of total RNA was used to generate the first strand of cDNA with the TUREscript cDNA Synthesize Kit (Aidlab Biotech Co. Ltd., Beijing).

### 2.3 Quantitative RT-PCR

The total RNA from the fat bodies of fifth-instar larvae was reverse transcribed into cDNAs. Real-time PCR was conducted with the specific primers F1 and R1 and F2 and R2 (Table 1) to determine the expression levels of *AP-sHSP20.8* and caspase-1, respectively. The housekeeping gene 18S rRNA (GenBank: DQ347469) was used to estimate equal amounts of RNA among the samples. Real-time PCR was performed in a StepOne Plus Real-Time PCR System using the SYBR^®^ Premix Ex Taq™ kit (TaKaRa). A 20 μL aliquot of reaction mixture contained 10 μL of 2× SYBR^®^ Premix Ex Taq™ buffer, 1 μL each of forward and reverse primers, 1 μL of cDNA, and 7 μL of RNase-free H2O. The PCR procedure was as follows: 95 °C for 10 s, followed by 40 cycles each at 95 °C for 15 s, at 62 °C for 15 s, and at 72 °C for 30 s. At the end of the reaction, as the sample was slowly heated from 60 to 95 °C, a melting curve was produced by monitoring the fluorescence continuously. Each independent experiment was conducted in triplicate, and the data were analysed using Student’s *t*-test. A *P* value <0.05 was statistically significant, and significance was indicated by an asterisk (Livak et al, 2001).

**Tab 1.**
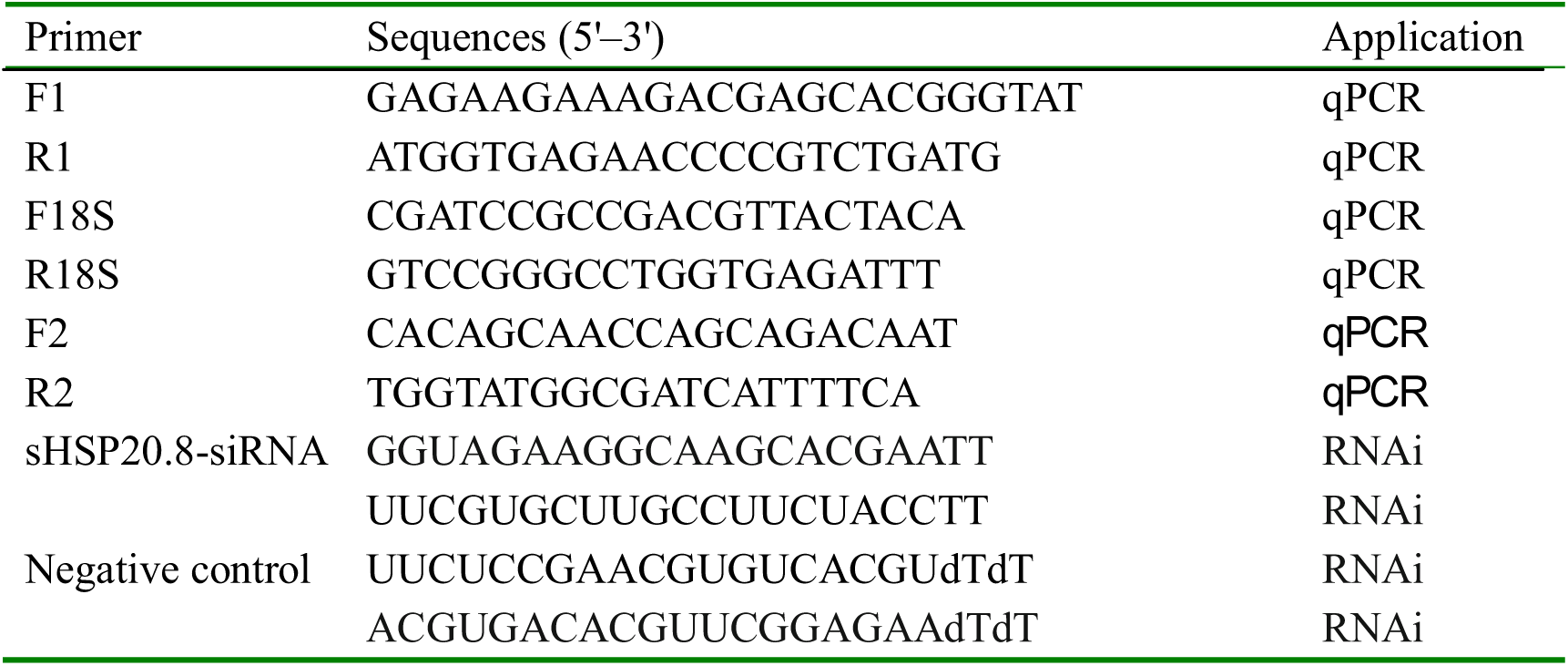
The primers used in this study

### 2.4. Western blotting

The samples of the fatbodies for proteins were ground in liquid nitrogen and dissolved in RIPA lysis buffer (Sangon, Shanghai, China). Protein concentrations in these extracts were determined using the Bradford method, and the samples were diluted to 2 μg/μL with phosphate-buffered saline (PBS).. After being separated by SDS-PAGE, the proteins were transferred to a polyvinylidene difluoride (PVDF) membrane by electrophoretic transfer. The membranes were blocked with 5% nonfat milk in PBST (PBS with 0.1% Tween-20) overnight at 4 °C and were then washed with PBST three times at 10 min intervals. The membranes were subsequently incubated with anti-Ap-sHSP20.8 antibody(Zhang et al,2014) which was diluted 1:500 with 5% nonfat milk in PBST, overnight at 4 °C. The membranes were washed with PBST and incubated with horseradish peroxidase (HRP)-conjugated goat anti-rabbit IgG (TransGen, Beijing, China), which was diluted 1:2000 with 5% nonfat milk in PBST, for 50 min at room temperature. Protein bands were detected with an HRP-DAB Detection Kit (Sangon, Shanghai, China).

### 2.5. RNA interference of Ap-sHSP20.8 gene

The siRNAs (Table 1) were designed by the siRNA Selection Program (<u>http://sirna.wi.mit.edu/home.php</u>) and were chemically synthesised by Shanghai GenePharma Co., Ltd. (Shanghai, China). The BLAST homology search (<u>http://www.ncbi.nlm.nih.gov/BLAST</u>) was performed to avoid off-target effects on other genes or sequences. The siRNAs were purified by high-performance liquid chromatography and were dissolved in diethylpyrocarbonate-treated water (Milli-Q-grade). The final concentration of siRNA was 1 μg/μL H_2_O. The 10 μL of siRNA was injected into each pupa using microliter syringes (Gaoge Co., Shanghai, China). To avoid leakage of siRNA from the insect body, needles were kept still at the injection point for 30 s. One set of siRNAs with random sequences was used as a negative control and was injected alongside the experimental injection. Twenty-three hours after RNAi treatment, the larvae were exposed to 43 °C for 0.5 h, followed by recovery at 25 °C for 0.5 h. The mixtures of the fatbodies were collected at 24 and 48 h after siRNA injection, frozen in liquid nitrogen and stored at -80 °C. All experiments were conducted with two independent experiments in triplicate.

### 2.6. Injection of insect and isolation of fat body

All inhibitors were purchased from Sigma Chemicals. After heat shock (HS) preconditioning (43 °C) for 30 min, A. pernyi was injected with RNAi or 10 μg of dexamethasone (DEX) to inhibit phospholipase A2, or 10 μg of indomethacin (Indo) or Naproxen (Nap) to inhibit cyclooxygenases, or 10 μg nordihydroguaiaretic acid (NDGA) or esculetin to inhibit lipoxygenase in 10 μL of DMSO, or DMSO alone (control). After the experiment, the fat bodies were excised, immediately frozen in liquid nitrogen and stored at -80 °C.

### 2.7. Caspase-1 activity assays

The activity of caspase-1 was determined with a Caspase-1 Activity Kit (Beyotime Institute of Biotechnology, Haimen, China), which was based on the ability of caspase-1 to change acetyl-Tyr-Val-Ala-Asp *p*-nitroanilide (Ac-YVAD-*p*NA) into the yellow formazan product *p*-nitroaniline (*p*NA). Cell lysates were centrifuged at 12,000 g for 10 min, and the protein concentrations were determined by the Bradford protein assay. The fat bodies extracts (30 μg of protein) were incubated in a 96-well microtiter plate with 20 ng of Ac-DEVD-*p*NA overnight at 37 °C. The absorbance values of *p*NA at 405 nm, OD405, were measured with a 96-well plate reader (BioTek, Santa Barbara, CA, USA). An increase in the OD405 indicated caspase-1 was activated.

## 3. Results

### 3.1. Effects of RNAi and eicosanoid biosynthesis inhibitors on the transcripts of Ap-sHSP20.8

The mRNA and protein abundance of Ap-sHSP20.8 were monitored at 24 and 48 h after siRNA injection and heat treatment. The Ap-sHSP20.8 gene and protein were successfully knocked down using sequence-specific si-sHSP20.8 RNA compared with the control group challenged with depc H2O at 24 and 48 h (Fig. 1A). However, the Ap-sHSP20.8 gene was successfully decreased in heat shock preconditioned larvae upon injection with eicosanoid biosynthesis inhibitors compared with the DMSO control (Fig. 1B). The small heat shock proteins(sHSPs) protect the activity of restriction enzymes and prevent thermal aggregation to retain their biological activity (Liu et al, 2013).

**Fig. 1.**
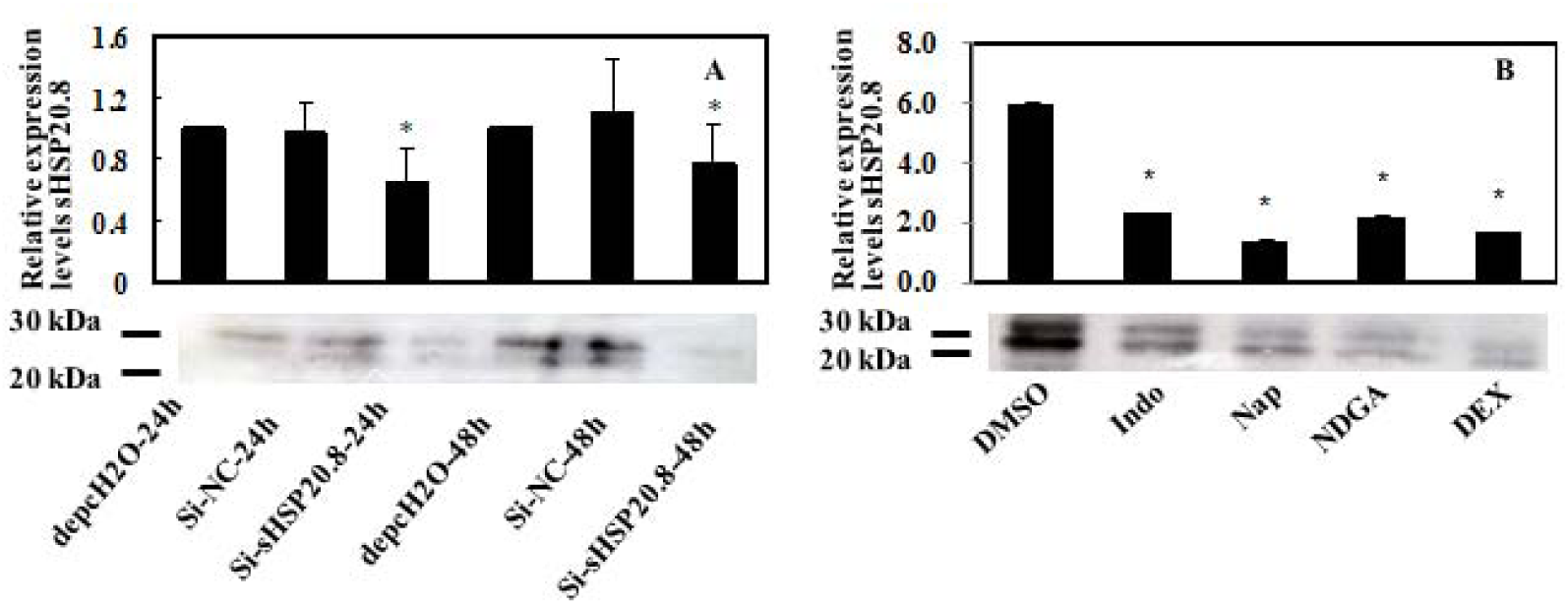
Relative mRNA expression levels and western blot analysis of *Ap-sHSP20.8* in response to the thermal stress by injection of siRNA at 24 and 48 h (A) and of eicosanoid biosynthesis inhibitors (B). A: Depc H2O and one set of negative control siRNAs (si-NC) were injected alongside the experimental injections. B: Larvae were first injected with 100 μ g (in 10 μ L DMSO) of either dexamethasone (DEX), nordihydroguaiaretic acid (NDGA) or indomethacin (INDO) or Naproxen(Nap). For control larvae, 10 μL DMSO was injected. After 30min they were treated at 43 °C for 0.5 h in a constant temperature incubator and then were kept at 25 °C for 1 h. After heat treatment, the fat bodies were immediately collected. Expression levels were assessed with the 18S rRNA gene for normalization. The data were analysed by Student’s *t*-test and presented as mean μ SE of independent experiments conducted in triplicate, and the asterisks represent the significant differences (p< 0.05, n = 5).

### 3.2. Expression of caspase-1 response to heat-si-sHSP20.8 and eicosanoid biosynthesis inhibitors

The transcripts of caspase-1 were significantly up-regulated with heat-si-sHSP20.8 (Fig. 2A), and the gene was also elevated when preconditioned with eicosanoid biosynthesis inhibitors (Fig. 2B). Furthermore, the maximal pharmacological effect was from Nap, which was an inhibitor of cyclooxygenases.

**Fig. 2.**
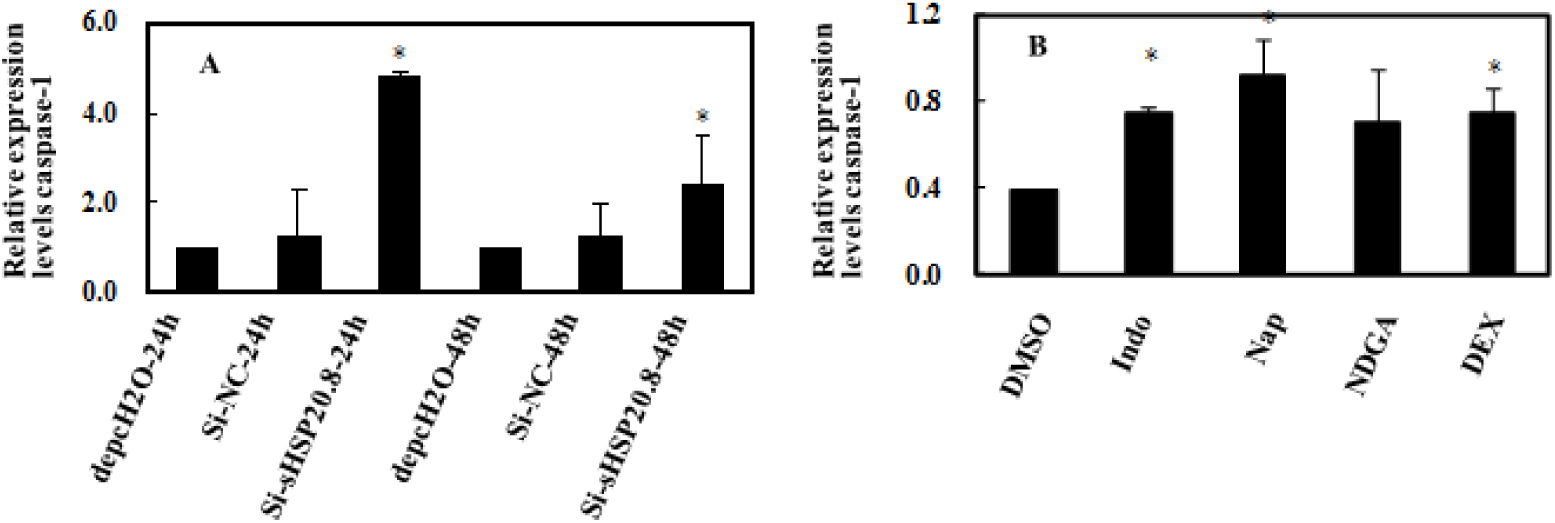
Different expression patterns of caspase-1 in response to the thermal stress by injection of siRNA at 24 and 48 h (A) and of eicosanoid biosynthesis inhibitors (B). A: Depc H2O and one set of negative control siRNAs (si-NC) were injected alongside the experimental injections. B: Larvae were first injected with 100 μg (in 10 μL DMSO) of either dexamethasone (DEX), nordihydroguaiaretic acid (NDGA) or indomethacin (INDO) or Naproxen(Nap).. For control larvae, 10 μL DMSO was injected. After 30min they were treated at 43 °C for 0.5 h in a constant temperature incubator and then were kept at 25 °C for 1 h. After heat treatment, the fat bodies were immediately collected. Expression levels were assessed with the 18S rRNA gene for normalization. The data were analysed by Student’s *t*-test and presented as mean μ SE of independent experiments conducted in triplicate, and the asterisks represent the significant differences (p< 0.05, n = 5).

### 3.3 Caspase-1 activity response to heat-si-sHSP20.8 and eicosanoid biosynthesis inhibitors

Preconditioning with si-sHSP20.8 RNAs activated caspase-1 activity upon subsequent heat shock, and this function was mediated by Hsp20.8 chaperone activity(Fig.3A). Exposure to any of the eicosanoid biosynthesis inhibitors was positively correlated with caspase-1 activity compared with control fat body preparations that were challenged with DMSO after heat shock(Fig.3B).

**Fig. 3.**
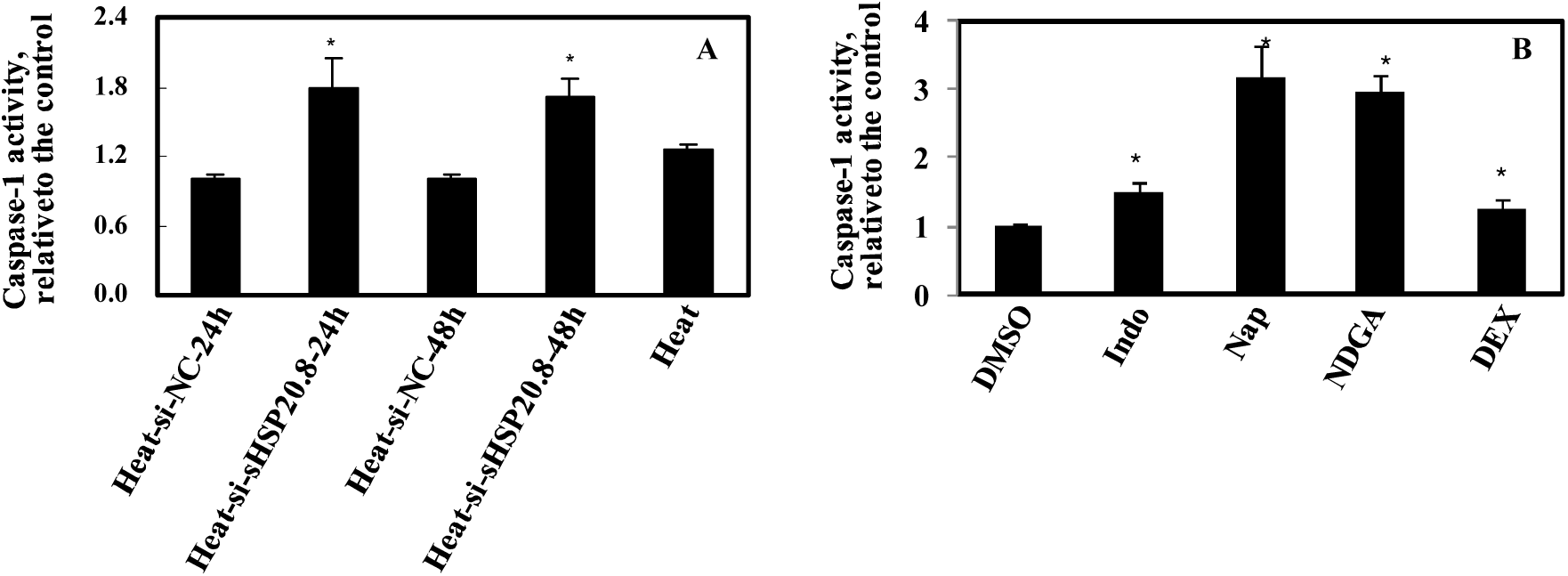
Effects on caspase-1 activities in response to heat shock in the fat bodies of *A.pernyi* by injection of siRNA at 24 and 48 h (A) and of eicosanoid biosynthesis inhibitors (B). The relative caspase-1 activities were calculated as ratios of the cleavage of their substrates in treated fat bodies relative to control, and the values of the control were set to 1. The data were analysed by Student’s *t*-test and presented as mean μ SE of independent experiments conducted in triplicate, and the asterisks represent the significant differences (p< 0.05, n = 5).

## 4. Discussion

The major finding of this study was that preconditioning with si-sHSP20.8 RNAs and eicosanoid biosynthesis inhibitors significantly decreased Ap-sHSP20.8 transcripts in the fat bodies of A. pernyi, whereas caspase-1 mRNA expression was simultaneously upregulated. Caspase-1 activity can also be pharmacologically modulated under such conditions. Additionally, protein expression of Ap-sHSP20.8 was also sensitive to inhibitors of the eicosanoid pathways and RNAi treatment. We suggested that the protocols that simultaneously up-regulated Hsp synthesis and downregulated caspase-1 activity could be used to increase the heat shock stress. We concluded that in the fat bodies of A. pernyi Hsp20.8 and eicosanoid biosynthesis inhibitors were negatively correlated with caspase-1 activity.

The heat shock response is a homeostatic adaptive response exhibited by nearly all cells in response to thermal and a variety of other stresses. This response is important because of the cytoprotective action of heat shock proteins during the pathogenesis of disease states, which includes changing temperatures, toxic substances, and particularly, various microbes(Zhou et al, 2009; Zhao and Jones, 2012; Shim et al, 2010). The cytoprotective effect of heat shock proteins appears to be mediated through several activities, including the dissociation of protein aggregates, a role in the folding of newly synthesised polypeptides(Arrigo, 2005; Liu et al, 2012), the modulation of the cellular detoxifying machinery to cope with oxidative stress(Creagh, 2000), and the modulation of cytoskeletal integrity(Mounier and Arrigo, 2002).

Apoptosis plays a role in mammalian development as a quality control mechanism to eliminate cells that are damaged, nonfunctional, abnormal, or misplaced(Zhuang et al, 2011; Stanley, 2011). The activation of caspases represents a critical step in the pathways that lead to the biochemical and morphological changes that underlie apoptosis. Previous studies showed that Hsp60 was able to substantially accelerate the maturation of procaspase-3 with different upstream activator caspases, and that this effect was dependent on ATP hydrolysis(Xanthoudakis et al,1999). Additionally, down-regulation or disruption of hsp70 expression resulted in apoptosis(Wei et al, 1995). In mammalian cells, the inhibition of SAPK/JNK signalling and apoptotic protease effector steps by hsp70 likely contributed to the resistance to stress-induced apoptosis observed in cells with transiently induced thermotolerance(Dick et al, 1997). Additionally, in *in vitro* systems, recombinant Hsp60 and Hsp10 accelerated the activation of procaspase-3 by cytochrome c and dATP in an ATP-dependent manner, which was consistent with their function as chaperones(Samali et al, 1999). In the present experiment, preconditioning with si-sHSP20.8 RNAs, the transcript level of caspase-1 upregulated approximately 5-fold compared with the control at 24 PI. The results indicated that sHSP might interrupt the activity of caspase-1 and defend against the heat-stress injury.

The inflammasome-dependent biosynthesis of eicosanoids is mediated by the activation of cytosolic phospholipase A2, which is specifically primed for the production of eicosanoids by the high expression of eicosanoid biosynthetic enzymes. Several possible mechanisms exist by which eicosanoid biosynthesis inhibitors could affect caspase-3 activity: the inhibition of AA release, the inhibition of leukotriene or PG synthesis, and the direct effect of AA downstream of the cytoprotective clusterin (Balogh et al,2013; Zhang et al, 2009). Considerable evidence now exists for the involvement of eicosanoids in the immune responses to bacteria, fungal, protozoan and parasitoid challenges in a phylogenetically wide range of insects. In *G. mellonella* and *Bombyx mori*, cyclooxygenase products, such as prostaglandins, mediated nodulation responses to viral infection(Durmus et al, 2008; Stanley-Samuelson et al, 1997). Similarly, bacterial challenge stimulated increased PLA2 activity in isolated haemocyte preparations of tobacco hornworms, *Manduca sexta*, relative to control haemocyte preparations that were challenged with water. Inflammasomes activated the caspase-1 protease, which processed the cytokines interleukin (IL)-1 beta and IL-18 and initiated a lytic host cell death called pyroptosis (von Moltke et al, 2012). In the present study, treating silkworms with the PLA2 inhibitor, dexamethasone, resulted in an approximately 6-fold increased expression level of *AP-sHSP20.8* compared with the control upon heat shock stress. The injection of cyclooxygenase inhibitors, Indo or Nap, and lipoxygenase inhibitors, NDGA, and caused 1.6-, 2.3-, 1.4- and 2.2-fold decreasing of *AP-sHSP20.8 gene transcripts*, respectively. However, the transcripts of caspase-1 were upregulated with DEX, Indo, Nap and NDGA were 0.25, 0.8, 0.6, 0.2 and 0.45-fold, respectively. Interestingly, the eicosanoid biosynthesis inhibitors stimulated the activity of caspase-1 to 1.5, 3.1, 2.9, and 1.25-fold of the control by DEX, Indo, Nap and NDGA compared with no heated group, respectively. All data suggested that cellular damages, including those induced by bacteria, virus and heat shock, might interrupt the biosynthesis of AA, and that caspase-1 might be directly involved in AA induced sHSP RNA expression. Together, the results in this paper help to extend the idea that eicosanoids mediate cellular immune responses in insects and to demonstrate the major elements of the eicosanoid system in silkworms. An increased understanding of insect adaptations to high temperatures will be important in the study of the evolution of insect insecticide resistance and for applications in integrated pest management.

## Acknowledgments

This work was supported by the earmarked fund for modern Argo-industry Technology Research Systems (CARS-22 SYZ10), the Anhui High Schools Natural Science Foundation (KJ2013B320), the National 863 plans projects of China (2011AA100306), the Sericulture Biotechnology Innovation Team (2013xkdt-05), and Ph.D. programs in Biochemistry and Molecular Biology (xk2013042).

